# Fast gene set enrichment analysis

**DOI:** 10.1101/060012

**Authors:** Gennady Korotkevich, Vladimir Sukhov, Nikolay Budin, Boris Shpak, Maxim N. Artyomov, Alexey Sergushichev

## Abstract

Gene set enrichment analysis (GSEA) is an ubiquitously used tool for evaluating pathway enrichment in transcriptional data. Typical experimental design consists in comparing two conditions with several replicates using a differential gene expression test followed by preranked GSEA performed against a collection of hundreds and thousands of pathways. However, the reference implementation of this method cannot accurately estimate small P-values, which significantly limits its sensitivity due to multiple hypotheses correction procedure.

Here we present FGSEA (Fast Gene Set Enrichment Analysis) method that is able to estimate arbitrarily low GSEA P-values with a high accuracy in a matter of minutes or even seconds. To confirm the accuracy of the method, we also developed an exact algorithm for GSEA P-values calculation for integer gene-level statistics. Using the exact algorithm as a reference we show that FGSEA is able to routinely estimate P-values up to 10^−100^ with a small and predictable estimation error. We systematically evaluate FGSEA on a collection of 605 datasets and show that FGSEA recovers much more statistically significant pathways compared to other implementations.

FGSEA is open source and available as an R package in Bioconductor (http://bioconductor.org/packages/fgsea/) and on GitHub (https://github.com/ctlab/fgsea/).

## 1 Main

Preranked gene set enrichment analysis (GSEA) [1] is a widely used method for analyzing gene expression data, particularly for datasets with small number of replicates. It allows to select from an *a priori* defined collection of pathways those which have non-random behavior in a considered experiment (Fig 1a). The method uses an enrichment score (ES) statistic which is calculated based on a vector of gene-level signed statistics, such as *t*-statistic from a differential expression test. As the analytical form of the null distribution for the ES statistic is not known, empirical null distribution has to be calculated. That can be done in a straightforward manner by sampling random gene sets as was done in the reference implementation [1] and reimplementations [2, 3], In this case for each of the input pathways a number of random gene sets of the same size are generated, and for each of them an ES value is calculated. Then a P-value is estimated as the number of random gene sets with the same or more extreme ES value divided by the total number of generated gene sets (a formal definition is available in the section 2.1). Finally, a multiple hypothesis correction procedure is applied to get adjusted P-values.

**Figure 1:**
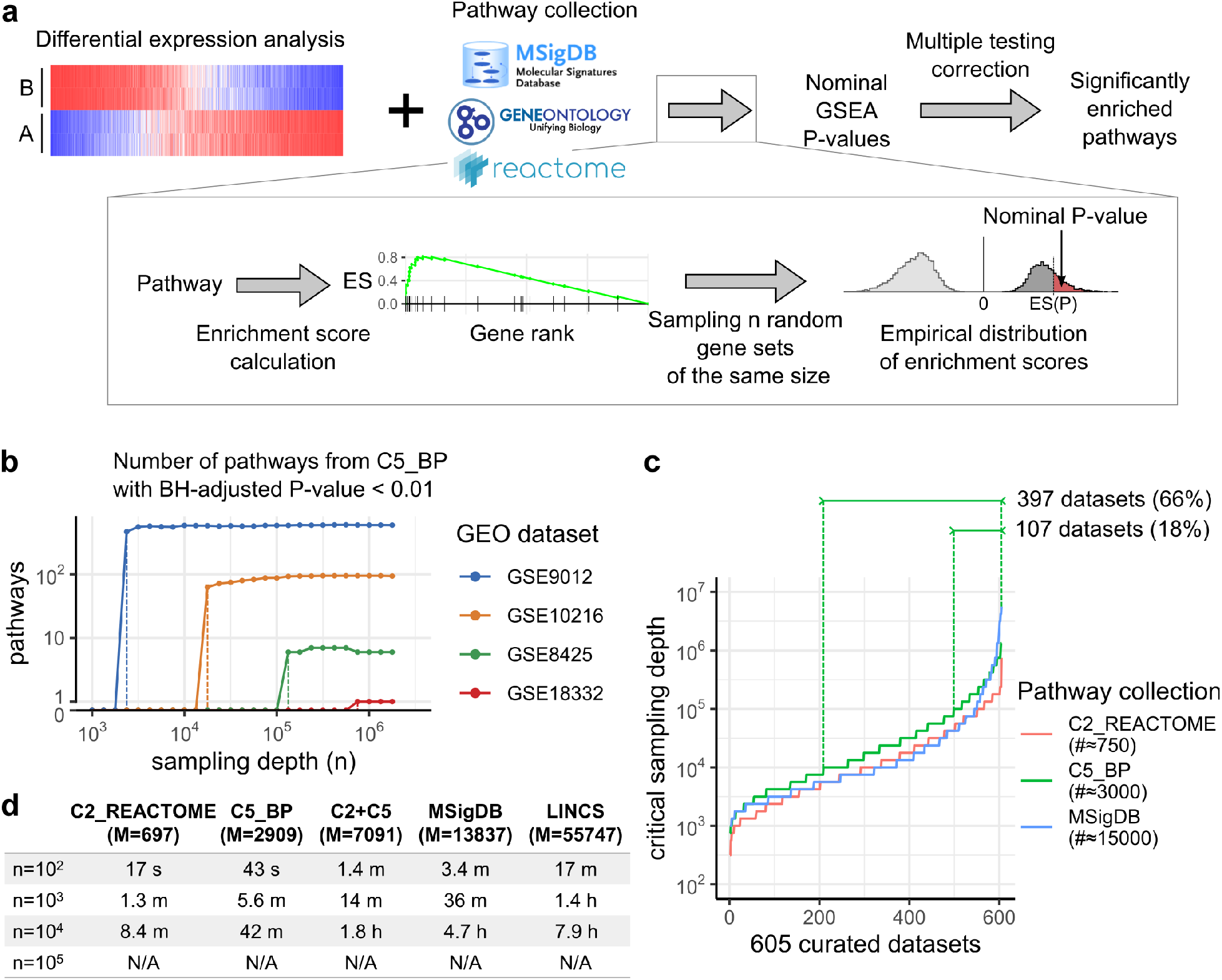
Gene Set Enrichment Analsyis (GSEA) sensitivity depends on the ability to reach high sampling depth, **a**, Overview of preranked GSEA method, **b**, The number of pathways reaching Benjamini-Hochberg adjusted P-value threshold depends on the sampling depth in a phase transition manner. The critical sampling depth, where the transition happens, varies with the dataset, **c**, Distribution of critical sampling depth values across 605 curated datasets from Gene Expression Omnibus, as estimated for three pathway collections, **d**, Time required for the reference GSEA implementation to estimate P-values for different gene set collections and sampling depth values for dataset GSE22293. N/A values indicate out of memory errors.

However, a large number of generated random gene sets can be required to reach a given false discovery rate (FDR) level on some datasets. As an example, we calculated GSEA P-values for Gene Ontology Biological Pathways collection (C5_BP subset of MSigDB collection [1]) on four datasets from Gene Expression Omnibus (GEO) varying the sampling depth, and calculated the number of pathways reaching FDR level of 0.01 after Benjamini-Hochberg (BH) correction. Due to the properties of BH procedure, the dependence of the number of significant pathways in these experiments has a phase transition behavior (Fig 1b): for each of the datasets there exist a certain critical sampling depth after which the number of significant pathways becomes non-zero and stays on the same level. This critical sampling depth is different for different datasets, but ultimately can reach the order of *M*/*α*, where *α* is the selected FDR threshold and *M* is the number of considered pathways.

To systematically asses the distribution of GSEA critical sampling depth on real datasets we prepared a collection of 605 microarray datasets from Gene Expression Omnibus (GEO) containing only two biological condistions. For each of these datasets we ran the differential expression analysis and used the results as gene-level statistics. We discovered that more than half of the datasets has the critical sampling depth of at least 10^4^ and a noticeable portion (10–20% depending on the collection) has the critical sampling depth of at least 10^5^ (Fig 1c). When a large pathway collection is considered (entire MSigDB collection) individual datasets has values of critical sampling depths reaching 5 · 10^6^. However, even running the reference implementation with the sampling depth of *n* = 10^4^ routinely is inconvenient and running it with *n* = 10^5^ can be impossible due to the time and memory consumption (Fig 1d): time and memory requirements grow linearly with the number of samples and the collection size.

To improve applicability of preranked GSEA analysis we present a fast gene set enrichment analysis (FGSEA) method for accurate and efficient estimation of GSEA P-values for a collection of pathways. The method consist of two main procedures: *FGSEA-simple* and *FGSEA-multilevel*. FGSEA-simple procedure allows to efficiently estimate P-values with a limited accuracy but simultaneously for the whole *collection* of gene sets, while FGSEA-multilevel procedure allows to accurately estimate arbitrarily low P-values but for *individual* gene sets.

FGSEA-simple procedure is based on an idea that generated random gene set samples can be shared between different input pathways. Indeed, consider *M* gene sets of the sizes *K*_1_ ⩽ *K*_2_ ⩽ … ⩽ *K_M_* = *K* and a collection of *n* independent samples *g_i_* of size *K* (Fig 2a). As in the naive approach, due to *g_i_* being independent samples of the size *K* the P-value for the pathway *M* can be estimated as a proportion of samples *g_i_* having the same or more extreme ES value as the pathway *M*. However, for any other pathway *j* we can construct a set of *n* independent samples of size *K_j_* by considering the prefixes *g*_*i*,1..*K_j_*_. Again, given a set of independent samples, the P-value can be estimated as a proportion of the samples having the same or more extreme ES value.

**Figure 2:**
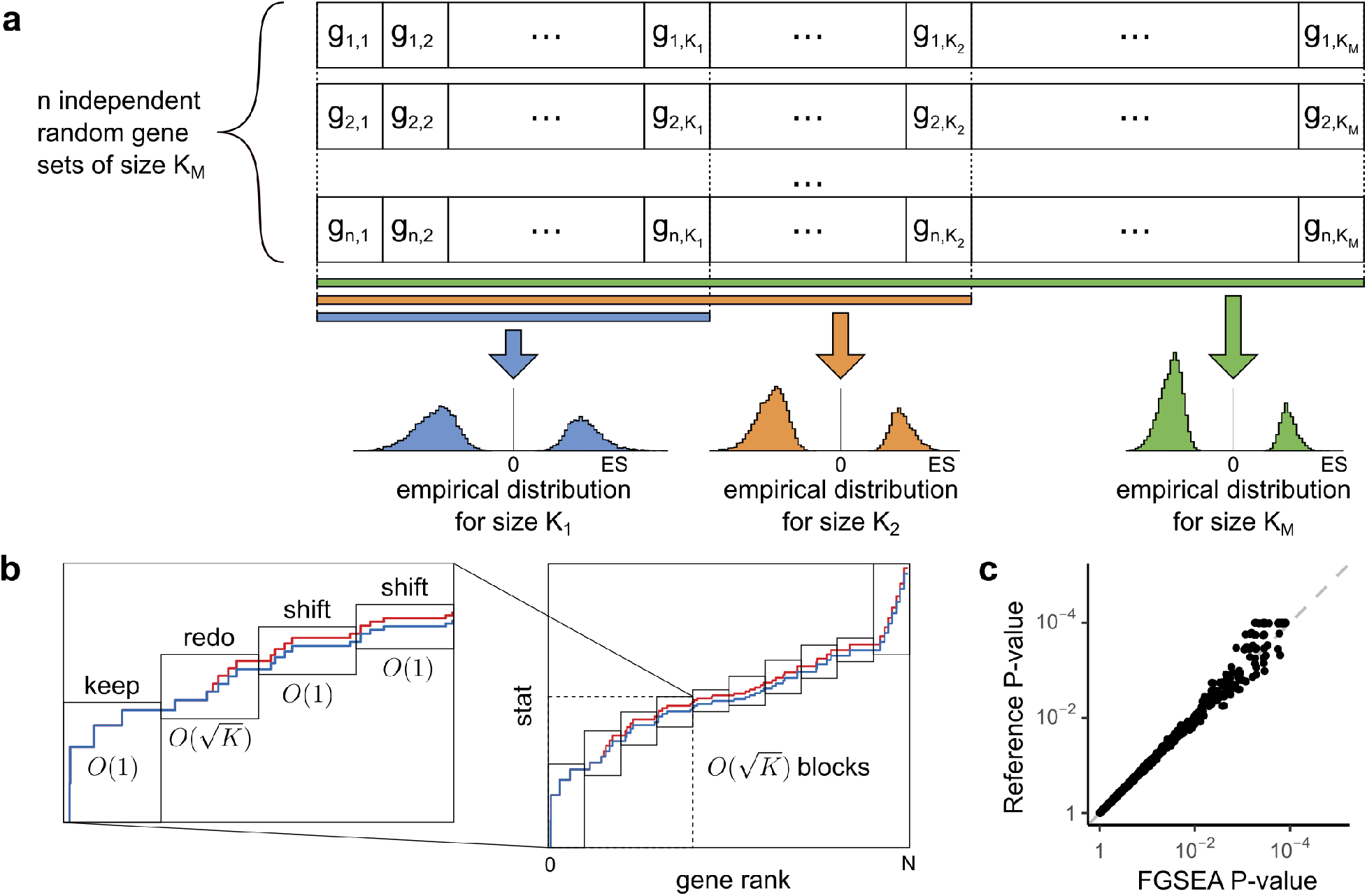
Preranked gene set enrichment analysis can be sped up by sharing sampling information between different gene set sizes, **a**, It is sufficient to generate *n* independent samples of size *K_M_* to calculate empirical distribution for any sizes *K_j_* ⩽ *K_M_* by considering only the prefix of the samples of the size *K_j_* **b**, For a given gene sample enrichment scores for all the prefixes can be efficiently calculated by employing a square root heuristic, **c**, The P-values calculated with the FGSEA-simple method are consistent with the reference implementation, but the results are obtained hundreds times faster.

The next important idea is that given a gene set sample *g_i_* of the size *K* the ES values for *all* the prefixes *g*_*i*,1..*j*_ can be calculated in an efficient manner using a square root heuristic (Fig 2b). Briefly, a variant of an enrichment curve is considered: the genes are enumerated starting from the most up-regulated to the most down-regulated, with the curve going to the right if the gene is not present in the pathway, and the curve goes upward if the gene is present in the pathway. It can be shown that the enrichment score can be easily calculated if curve point most distant from the diagonal is known. Let us split *K* genes from the gene set into 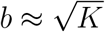 consecutive blocks of size 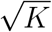 and consider what happens with the curve when we change the prefix from *g*_*i*,1..*j*−1_ to *g*_*i*,1..*j*_ by adding gene *g_i,j_*. The curve in the blocks to the left of *g_i,j_* are not changed at all, while the blocks to the right of *g_i,j_* are uniformly shifted. This observation allows us to consider the prefixes in an increasing order and update the position of the most distant point in 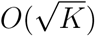 time. Briefly, for the each block which is either not changed or shifted the update procedure takes *O*(1) time, while for the changed block the update procedure is proportional to its size and takes 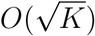 time. Finally, aggregating the blocks takes additional 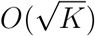 time. Overall this results in time complexity of 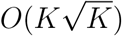 to calculate ES values for all the prefixes. In total, the time complexity of the calculating P-values for the set of *M* pathways is 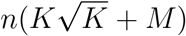, which gives around 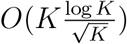 speed up compared to a naive approach. The full description of the algorithm is given in the section 2.3.

As an example we ran FGSEA-simple and the reference implementations on the same example dataset of genes differentially regulated on Th1 activation [4] against a set of 700 Reactome [5] pathways (see section 2.2) and compared the resulting nominal P-values (Fig 2c). Both methods were ran with *n* = 10000 and the results are indistinguishable from each other up to the random noise inherent to both methods. However, on this example the reference implementation took about 420 seconds, while FGSEA-simple finished in about 4 seconds. The two order of magnitude speed-up is consistent with the theoretical one due to the algorithm time complexity. Given a highly parallel implementation of FGSEA-simple, its performance allows to routinely achieve sampling depth of 10^5^ and accurately estimate P-values as low as 10^−5^.

However, accurately estimating P-values lower than 10^−5^ with FGSEA-simple can be impractical or even infeasible. To estimate such low P-values we developed *FGSEA-multilevel* method, which is based on an adaptive multi-level split Monte Carlo scheme [6], The method takes as an input an ES value *γ* > 0 and a gene set size *K*, and calculates the probability *P_K_* (ES ⩾ *γ*) of a random gene set of size *K* to have an enrichment score no less than *γ*. The method sequentially finds ES levels *l_i_* for which the probability *P_K_*(ES ⩾ *l_i_*) is approximately equal to 2^−*i*^ (see Fig 3a for a toy example). The method stops when *l_i_* becomes greater than *γ* and the P-value can be crudely approximated as 2^−*i*^.

**Figure 3:**
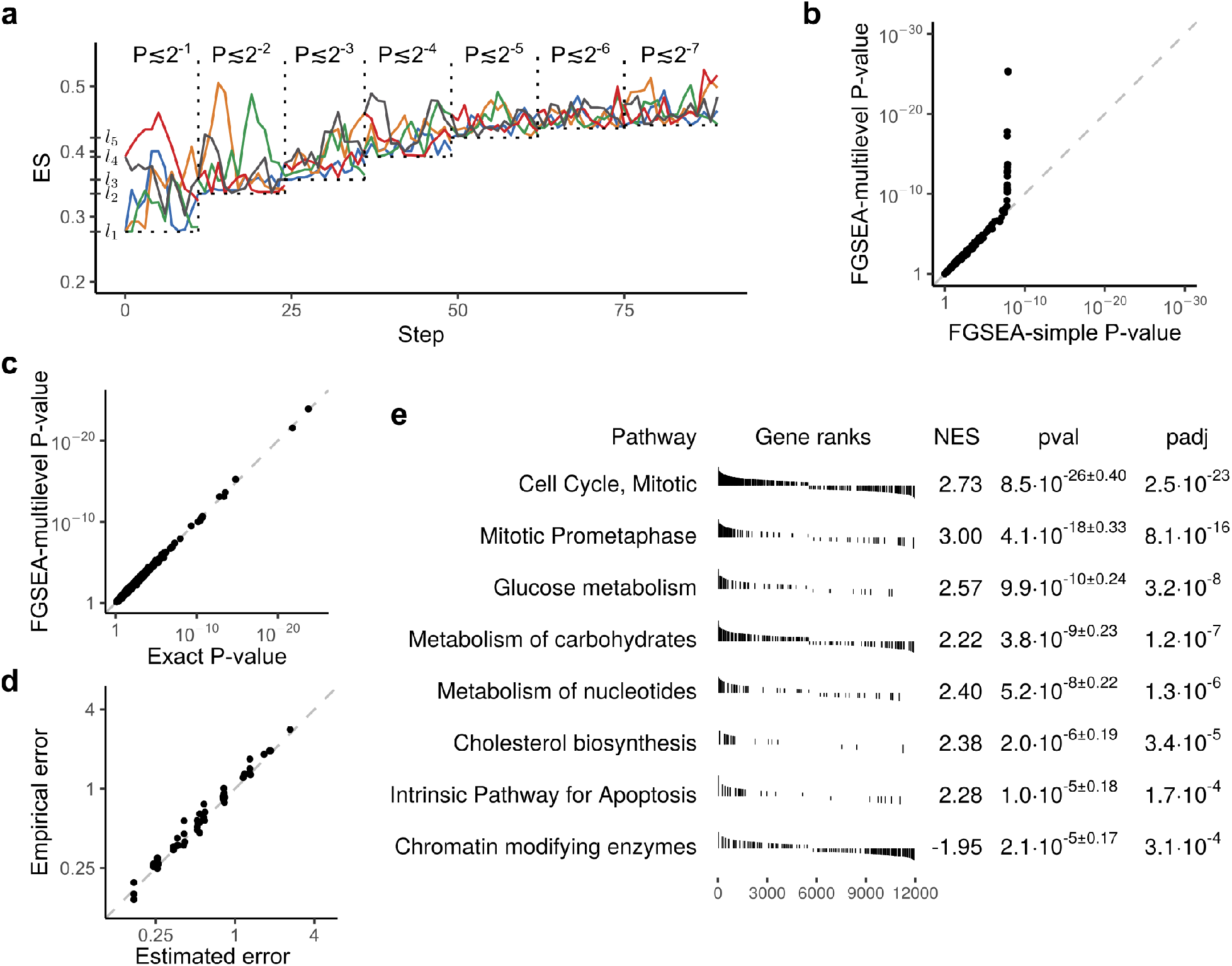
Adaptive multilevel Monte Carlo sampling scheme can be used to calculate arbitrarily low P-values. **a**, A toy illustration of the multilevel split Monte Carlo scheme for sample size of *Z* = 5. **b**, Comparison of GSEA P-values as calculated by FGSEA-simple method run with the sampleing depth *n* = 10^8^ and FGSEA-multilevel with the sample size of Z=101. **c**, Comparison of P-values as calculated with an exact method and FGSEA-multilevel method. Both methods were run on gene-level statistic values rounded to integers. **d**, Comparison between estimated and an observed error of *log*_2_ P-values for different P-values (from 10^−4^ to 10^−100^), gene set sizes (from 15 to 250) and sample sizes (from 101 to 1001). **d**, An example of FGSEA results as run with FGSEA-multilevel method for Th0 vs Th1 comparison and Reactome pathways. The analysis was run with sample size of *Z* =101. Redundant pathways were filtered.

The intermediate *l_i_* thresholds are calculated as follows. First, a set of *Z* (an odd number, parameter of the method) random gene sets of size *K* are generated uniformly and ES values for them are calculated. The median value of the ES values is calculated and assigned to *l*_1_. By construction, the probability *P_K_*(*γ* ⩾ *l*_1_) of a random gene set to have an ES value no less than *l*_1_ can be approximated as 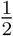. Next 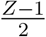 generated gene sets with the ES values less than *l*_1_ are discarded, while 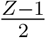 gene sets with the ES values greater than *l*_1_ are duplicated. This results in a sample of *Z* gene sets with the ES values no less than *l*_1_, but the distribution is non-uniform. However, it can be made into a uniform sample with a Metropolis algorithm. On each Metropolis algorithm step each gene set sample is tried to be modified by swapping a random gene from the set with a gene outside of the set. The change is accepted if an enrichment score of the new set is no less then current threshold *l*_1_, otherwise the change is rejected. Metropolis algorithm guarantees, that after enough steps the sample becomes close to uniformly distributed. Thus, a median of the enrichment scores (*l*_2_) would correspond to probability of 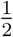 for a gene set to have an enrichment score no less than *l*_2_ given it has an enrichment score no less than *l*_1_:

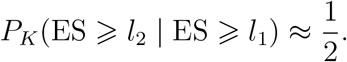

Which means

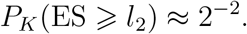

The same procedure is applied to calculate the next *l_i_* values.

The iterations stop when *l_i_* becomes greater than *γ*. On this iteration the probability of a random gene set to have a ES value no less than *γ* can be approximated as:

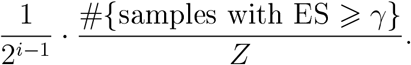

When estimating small P-values it becomes practical to carry out the estimation in logscale. In particular, the values become practically unbiased both in median and mean sense and it becomes simple to estimate the approximation error and condifence intervals (see section 2.5.4).

The full formal description of the algorithm is available in the section 2.5.

For the example dataset we show that P-values are as low as 10^−26^ for some of the pathways and the results are consistent with FGSEA-simple P-values ran on 10^8^ permutations (Fig 3b). Note, that FGSEA-multilevel calculation with sample size of Z=101 took only 10 seconds working on a single thread while 10^8^ permutations on FGSEA-simple took 40 minutes working in 32 threads.

To further prove the approximation quality of FGSEA-multilevel algorithm we developed an exact method for calculating GSEA P-values, but limited to integer weights. The method is based on dynamic programming, the full description is given in section 2.4. The complexity of the algorithm is *O*(*NKT*^2^), where *N* is the number of genes, *K* is the size of gene sets and *T* is the sum of the top *K* absolute values of gene-level statistics. With a number of optimizations this method allows to calculate P-values for rounded weights in the example dataset in a couple of hours.

When run on the same integer weights FGSEA-multilevel and the exact method give highly concordant results (Fig 3c). Additionally, using the exact P-values, real approximation errors can be compared with the estimated ones. We show, that the FGSEA-multilevel error estimation are highly concordant with the real errors (Fig 3d) for a wide range of P-values (from 10^−4^ to 10^−100^), gene set sizes (from 15 to 250) and sample sizes (from 101 to 1001).

In practice FGSEA-multilevel method is combined with FGSEA-simple. First, for all the input pathways FGSEA-simple method can be run with a limited sample size. Next, for the pathways that have high relative error after FGSEA-simple (i.e. pathways with low p-values) FGSEA-multilevel method is executed. As many of the pathways in an input collection usually are not enriched, they have a relatively high P-value and will be batch-processed with a highly efficient FGSEA-simple algorithm with deterministic time boundaries. The more interesting pathways with lower P-values will then be processed with FGSEA-multilevel algorithm individually and the amount of processing time will depend on their P-values.

As FGSEA allows to practically estimate the P-values for a large collections of gene sets, it can lead to a large number of statistically significant hits with high overlaps. To deal with this issue and make the representation of FGSEA results more concise we developed a procedure to filter the redundant gene sets. The procedure is similar to GO Trimming method [7] but is based on the Bayesian network construction approaches. It considers the significant pathways one by one and tries to remove gene sets that do not provide new information given some other pathway already present in the output. In this case, we consider a pathway *P*_1_ to give a new information given a pathway *P*_2_ if the P-value of pathway *P*_1_ in the universe of genes just from *P*_2_, or just from genes outside of *P*_2_, is less than some threshold. This procedure allows to filter redundant pathways without requirement of having any explicit hierarchy of pathways. The full description of the procedure is given in section 2.6. The table resulting from running FGSEA on the example dataset with filtering of redundant hits is shown on Fig 3e.

Finally, we have explored FGSEA performance on the collection of 605 curated GEO datasets described earlier and the C5_BP pathway collection. Notably, it took less then a minute per dataset of running time to finish FGSEA analysis with the multilevel algorithm (Fig 4a) on a laptop with 4-core Intel Core i5 processor, with a median time of 8 seconds. Besides the multilevel algorithm, the same analysis was carried out with FGSEA-simple algorithm with the sampling depth values of *n* = 10/ 10^4^ and 10^5^, and the reference implementation (Broad GSEA) with the sampling depth of 10^4^. For all these methods we compared the number of pathways reaching FDR level of 0.01: BH-adjusted P-values were used for FGSEA and reported Q-values (*Broad Q-values*) were used for the reference implementation (Fig 4b). The results reiterate that even sampling depth of 10^5^ is not enough to detect statistically significant enriched pathways for some of the datasets when BH-adjustment procedure is used.

**Figure 4:**
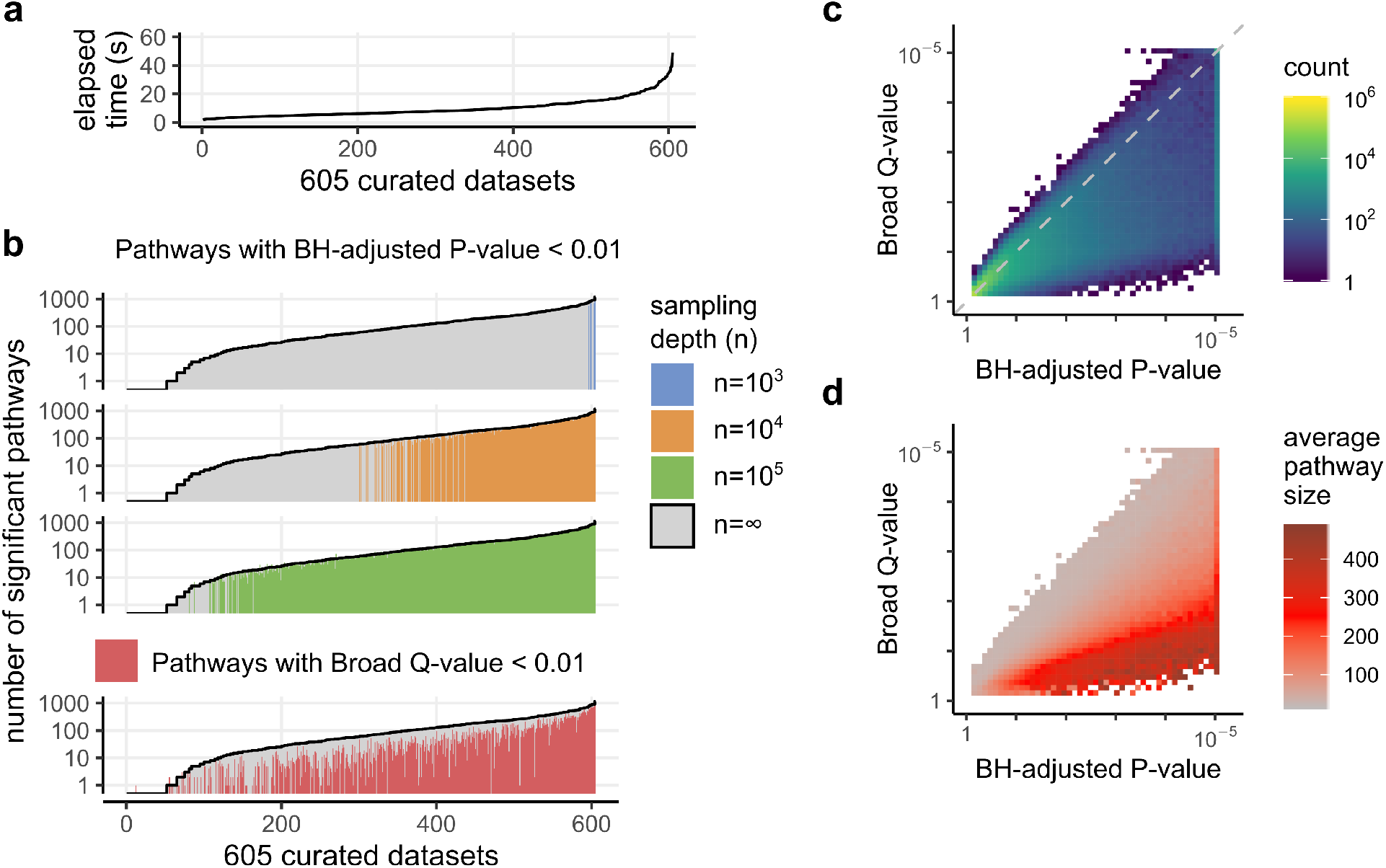
FGSEA method with the multilevel approach detects more statistically significant pathways compared to other GSEA implementations in a fraction of time. All plots represent data from analysis of the collection of 605 cureated GEO datasets with C5_BP used as a gene set collection, **a**, Wall clock time of running FGSEA. **b**, The number of detected statistically significant pathways for different adjusted P-value calculation procedures. Pathways with small BH-adjusted P-value after FGSEA-multilevel are shown in grey. Pathways with small BH-adjusted P-value after FGSEA-simple are shown in blue, orange and green, depending on the sampling depth. Pathways with small Q-values as reported by Broad GSEA are shown in red. **c**, Comparison of BH-adjusted P-values for FGSEA-multilevel and Q-values reported by Broad GSEA. For illustration purposes all values are capped at 10^−5^. **d**, Same as *c* but average pathway sizes are shown.

The *ad hoc* procedure implemented in Broad GSEA aggregates ES values generated across different pathways increasing the sensitivity on some of the datasets compared to BH-adjusted P-values for the same sampling depth of 10^4^, however this increase in sensitivity comes with overall more conservative behavior. The total number of pathways reaching FDR level of 0.01 for Broad Q-values is 39467, which is only 60% of 65690 pathways for BH-adjustment P-values with the sampling depth of 10^4^ and 48% of 81628 pathways for the multilevel algorithm. We further characterized this behavior by directly comparing Broad Q-values with BH-adjusted P-values and have shown that Broad Q-values are individually more conservative (Fig 4c) in a pathway size dependent manner (Fig 4d).

To conclude, here we have presented FGSEA method for fast preranked gene set enrichment analysis. The method allows to routinely estimate even very low P-values and can be used with conjunction with standard multiple hypothesis testing correction methods, such as Benjamini-Hochberg procedure. This, in turn, leads to better sensitivity and the ability to detect significant pathways in hard cases, where other implementations fail. FGSEA method is freely available as an R package at Bioconductor (http://bioconductor.org/packages/fgsea) and on GitHub (https://github.com/ctlab/fgsea).

## 2 Methods

### 2.1 Formal definitions

The preranked gene set enrichment analysis takes as input two objects: an array of gene-level statistic values *S* for the genes *U* = {1, 2,…, *N*} and a list of query gene sets (pathways) *P*. The goal of the analysis is to determine which of the gene sets from *P* has a non-random behavior.

The statistic array *S* of the size |*S*| = *N* for each gene *i* ∈ *U* contains a value 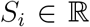 that characterizes the gene behavior in a considered biological process. Commonly, if *S_i_* > 0 the expression of gene *i* goes up on treatment compared to control and *S_i_* < 0 means that the expression goes down. Absolute values |*S_i_*| represent magnitude of the change. Array *S* is sorted in a decreasing order: *S_i_* > *S_j_* for *i* < *j*. The value of *N* in practice is about 10000–20000.

The list of gene sets *P* = {*P*_1_, *P*_2_,…, *P_M_*} of length *M* usually contains groups of genes that are commonly regulated in some biological process. We assume that the gene sets *P_i_* are ordered by their size (denoted as *K_i_*): *K*_1_ ⩽ *K*_2_ ⩽ … ⩽ *K_M_* = *K*. Usually only relatively small gene sets are considered with *K* ≈ 500 genes.

To quantify a co-regulation of genes in a gene set *p* Subramanian *et al*.[1] introduced a gene set enrichment score function *s_r_*(*p*) that uses gene rankings (values of *S*). The more positive is the value of *s_r_*(*p*) the more enriched the gene set is in the positively-regulated genes (with *S_i_* > 0). Accordingly, negative *s_r_*(*p*) corresponds to enrichment in the negatively regulated genes.

Value of *s_r_*(*p*) can be calculated as follows. Let *k* = |*p*|, NS = Σ_*i*∈*p*_|*S_i_*|. Let al so ES be an array specified by the following formula:

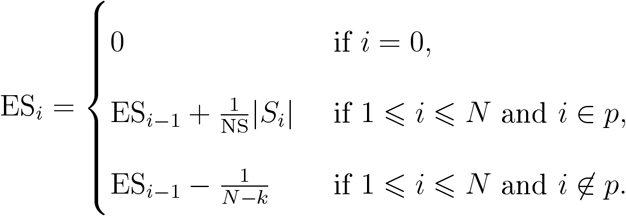

The value of *s_r_*(*p*) corresponds to the largest by the absolute value entry of ES:

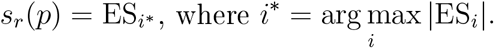

For convenience, we also introduce the following notation:

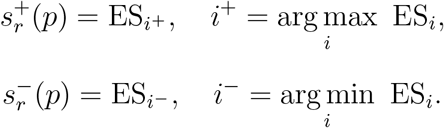

From these two values it easy to find value of *s_r_*(*p*), which is equal to 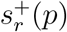 if 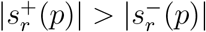 or 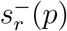 otherwise.

Often we will consider only the positive values of the gene set enrichment score function since:

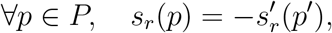

where 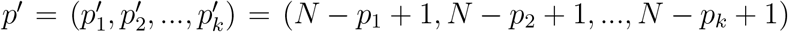 and 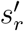 corresponds to the gene set enrichment score function for array *S*′ such that 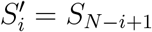.

Next, following Subramanian *et al* for a pathway *p* we define GSEA P-value as:

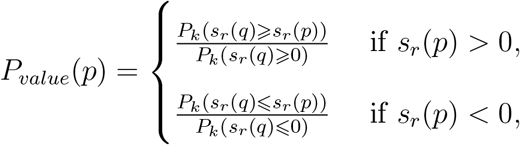

where *q* is a random gene set of size *k*.

### 2.2 The datasets

A collection of 605 curated datasets was generated from a set of all curated datasets (GDS) in Gene Expression Omnibus. Only the datasets with two biological conditions were kept. Differential expression was done using limma. The moderated t-statistic was used for gene ranking. Only top 10000 genes by average expression were used in ranking. The final rankings are available at https://ctlab.itmo.ru/files/software/fgsea/geo_ranks/.

Pathway collections C2_REACTOME, C5_BP, C2, C5, MSigDB were obtained via msigdbr package. LINCS perturbation collection was downloaded from Enrichr web-site. The corresponding gmt files are available at https://ctlab.itmo.ru/files/software/fgsea/gmts/.

As the example ranking ThO vs Thl comparison was used from dataset GSE14308 [4], The differential expression was calculated using limma [8], Only top 12000 genes by mean expression were used. Limma t-statistic was used as gene-level statistic. The script to generate rankings is available on GitHub: https://github.com/ctlab/fgsea/blob/master/inst/gen_gene_ranks.R.

Reactome [5] database was used as an example collection via reactome.db R package. For the analysis only the pathways of the size from 15 to 500 were used. The script to generate pathway collection is available on GitHub: https://github.com/ctlab/fgsea/blob/master/inst/gene_reactome_pathwavs.R

### 2.3 FGSEA-simple: an algorithm for fast calculation of GSEA P-values simultaneously for many pathways

In this section we describe an algorithm for fast estimation of GSEA P-values simultaneously for a collection of pathways *P*. There, for each pathway *p* a set of *n* uniformly random gene sets *q_i_* are considered. Then P-value is estimated as:

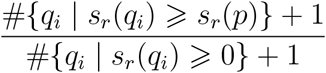

for positively enriched pathway *p* and as:

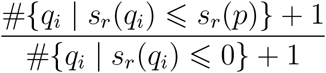

for negatively enriched pathway. These two formulas follow Subramanian *et al*. implementation, except of +1 terms, which are recommended by Phipson and Smyth [9], Otherwise, the nominal P-values from FGSEA-simple and reference implementation are indistinguishable, however FGSEA-simple works orders of magnitude faster.

#### 2.3.1 Cumulative statistic calculation for the mean statistic

Let first describe the idea of the proposed algorithm on a simple mean statistic *s_m_*:

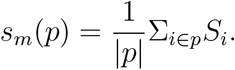

The main idea of the algorithm is to reuse sampling for different query gene sets. This can be done due to the fact that for an estimation of null distributions samples have to be independent only for a specific gene set size, while they can be dependent between different sizes.

Instead of generating *nM* independent random gene sets: *n* for each of *M* input gene sets, we will generate only *n* random gene sets of size *K*. Let *π_i_* be an *i*-th random gene set of size *K*. From that gene set we can generate gene sets for a all the query pathways *P_j_* by using its prefix: *π_i,j_* = *π_i_*[1..*K_j_*].

The next step is to calculate the enrichment scores for all gene sets *π_i,j_*. Instead of calculating enrichment scores separately for each gene set we will calculate simultaneously scores for all *π_i,j_* for a fixed *i*. Using a simple procedure it can be done in Θ(*K*) time.

Let us find enrichment scores for all prefixes of *π_i_*. This can be done by element-wise dividing of cumulative sums array by the length of the corresponding prefix:

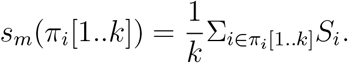

Selecting only the required prefixes takes an additional Θ(*m*) time.

The described procedure allows to find P-values for all query gene sets in Θ(*n*(*K* + *m*)) time. This is about min(*K, m*) times faster than the straightforward procedure.

#### 2.3.2 Cumulative statistic calculation for enrichment score

For the enrichment score *S_r_* we use the similar idea as above: we will also be sampling only gene sets of size *K* and from that sample will calculate statistic values for all the other sizes. However, calculation of the cumulative statistic values for the subsamples is more complex in this case. In this section we only be considering the positive mode of enrichment statistic 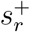.

It is helpful to look at enrichment score from a geometric point of view. Let us consider for a pathway *p* of size |*p*| = *k* a graph of *N* + 1 points (Fig. S1) with the coordinates (*x_i_, y_i_*) for 0 ⩽ *i* ⩽ *N* such that:

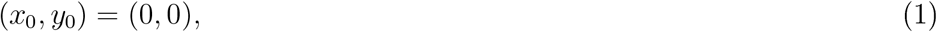

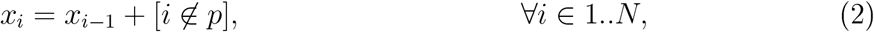

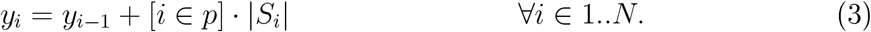

The calculation of 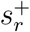 corresponds to finding the point farthest up from a diagonal ((*x*_0_, *y*_0_), (*x_N_, y_N_*)). Indeed, it is easy to see that *x_N_* = *N* – |*p*| = *N* – *K* and *y_N_* = Σ_*j*∈*p*_|*S_j_*| = NS, while the individual enrichment scores ES_*i*_ can be calculated as 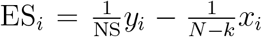. Value of ES_*i*_ is proportional to the directed distance from the line going through (*x*_0_, *y*_0_) and (*x_N_, y_N_*) to the point (*x_i_, y_i_*).

Let us fix a sample *π* of size *K*. To efficiently calculate cumulative values 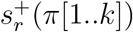 for all *k* ⩽ *K* we need a fast method of updating the farthest point when a new gene is added. In that case we can add genes from *π* one by one and calculate values 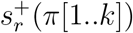 from the corresponding maximal distances.

Because we are calculating values for *π*[1..*k*] for *k* ⩽ *K* we know in advance which *K* genes will be added. This allows us to consider *K* + 1 points instead of *N* + 1 for each iteration *k*. Let array *o* of size *K* contain the sorted order of genes in *π*: that is, *π*_*o*_1__ is the minimal among *π, π*_*θ*_2__ is the second minimal and so on. The coordinates can be calculated as follows:

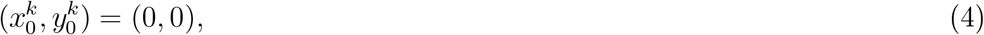

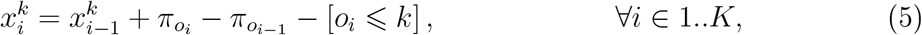

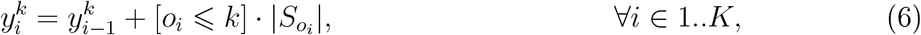

where we set *π*_*θ*_0__ to be zero.

It can be shown that finding the farthest up point among (4)–(6) is equivalent to finding the farthest up point among (1)–(3) with 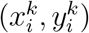 being equal to 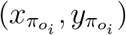 calculated for *p* = *π*[1..*k*]. Consider 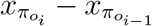. By the definition of *x* it is equal to:

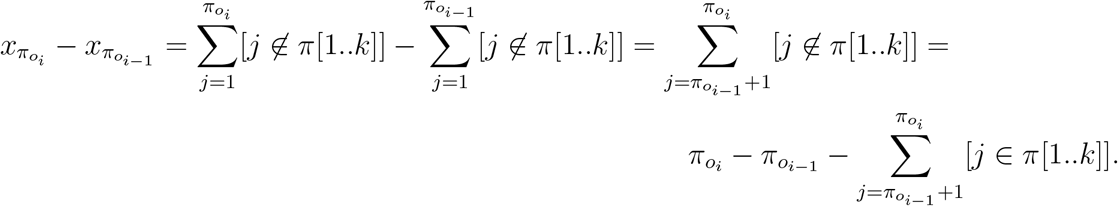

By the definition of *o*, in the interval [*π*_*o*_*i*-1__ + 1, *π_o_i__* – 1] there are no genes from *π* and, thus, from *π*[1..*k*]. Thus we can replace the sum with its last member:

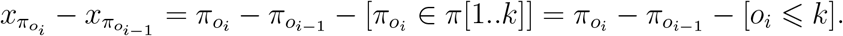

We got the same difference as in (5).

Now consider 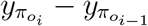. By the definition of *y* it is equal to:

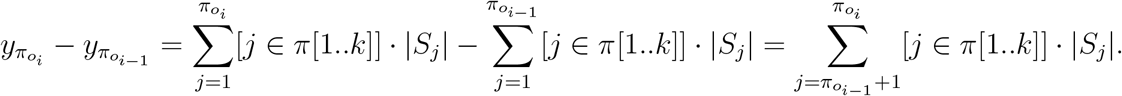

Again, in the interval [*π*_*o*_*i*−1__ + 1, *π_o_i__* – 1] there are no genes from *π*[1..*k*]. Thus we can replace the sum with only the last member:

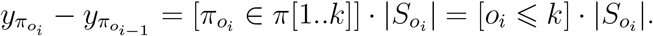

We got the same difference as in (6).

We do not need to consider other points, because points from *o*_*i*−1_ to *o_i_* − 1 have the same *y* coordinate and *o*_*i*−1_ is the leftmost of them. Thus, when at least one gene is added the diagonal ((*x*_0_, *y*_0_), (*x_N_, y_N_*)) is not horizontal and *o*_*i*−1_ is the farthest point among *o*_*i*−1_,…, *o_i_* − 1.

Now let consider what happens with the enrichment score graph when gene *π_k_* is added to the query set *π*[1..*k* – 1] (Fig. S2). Let *r_k_* be a rank of gene *π*_2_ among genes *π*, then coordinate of points (*x_i_, y_i_*) for *i* < *r_k_* do not change, while all (*x_i_, y_i_*) for *i* ⩾ *r_k_* are changed on (Δ_*x*_, Δ_*y*_) = (−1, |*S_π_k__*|).

To make fast incremental updates we will decompose the problem into multiple smaller ones. For simplicity we assume that *K* + 1 is an exact square of an integer *b*. Let split *K* +1 points into *b* consecutive blocks of the size *b*: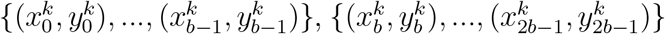 and so on.

For each of *b* blocks we will store and update the farthest up point from the diagonal. When we know for each block its farthest point we can find the globally farthest point by a simple pass in *O*(*b*) time.

Next, we show how to update the farthest points in blocks in amortized time *O*(*b*). This taken together with one *O*(*b*) pass will get us an algorithm to update the globally farthest point in amortized *O*(*b*) time.

Below we use *c* = ⌊*r_k_*/*b*⌋ as an index of a block where gene *π_k_* belongs, where *r_k_* is the ranking of the genes from *π*, i.e. *r_o_i__* = *i*.

First, we describe the procedure to update point coordinates. We will store *x_i_* coordinates using two vectors: *B* of size *b* and D of size *K* + 1, such th at *x_i_* = *B_i/b_* + *D_i_*. When gene *π_k_* is added all *x_i_* for *i* ⩾ *r_k_* are decremented by one. To reflect this we will decrement all *B_j_* for *j* > *c* and decrement all *D_i_* for *r_k_* ⩽ *i* < *cb*. The update takes *O*(*b*) time. After this update procedure we can get value *x_i_* in *O*(1) time. The same procedure is applied for *y* coordinates.

Second, for each block we will maintain an upper part of its convex hull. Having convex hull is useful because the farthest point in block always lays on its convex hull. All blocks except *c* have the points either not changed or shifted simultaneously on the same value. That means that the lists of points on the convex hulls for these blocks remain unchanged. For the block *c* we can reconstruct convex hull from scratch using Graham scan algorithm [10].

Because the points are already sorted by *x* coordinate, this reconstruction takes *O*(*b*) time. In total, it takes *O*(*b*) time to update the convex hulls.

Third, the farthest points in blocks can be updated using the stored convex hulls. Consider a block where the convex hull was not changed (every block except, possibly, block *c*). Because diagonal always rotates in the same counterclockwise direction, the farthest point in block on iteration *k* either stays the same or moves on the convex hull to the left of the farthest point on the (*k* − 1)-th iteration. Thus, for each such block we can compare current farthest point with its left neighbor on the convex hull and update the point if necessary. It is repeated until the next neighbor is closer to the diagonal than the current farthest point. In the block c we just find the farthest point in a single pass by the points on the convex hull.

To show that the updating the farthest points takes *O*(*b*) amortized time we will use potential method. Let a potential after adding *k*-th gene Φ_*k*_ be a sum of relative indexes of the farthest points for all the blocks. As there are *b* blocks of size *b* the sum of relative indexes lies between 0 and *b*^2^. Thus, Φ_*k*_ = *O*(*b*^2^). For an update of all *b* − 1 blocks except *c* we need to make *t_k_* = *b* − 1 + *z* operations of comparing two points, where *z* is the number of times the farthest points were updated. This can take up to Θ(*b*^2^) time in the worst case. However, it can be noticed, that potential change Φ_*k*_ − Φ_*k*−1_ is equal to −*z* + *O*(*b*): the sum of indexes is decreased by a number of times the farthest points were updated plus *O*(*b*) for the block c where the index can go from 0 to *b* − 1. This gives an amortized cost of *k*-th iteration to be *a_k_* = *t_k_* + Φ_*k*_ − Φ_*k*−1_ = *b* − 1 + *z* − *z* + *O*(*b*) = *O*(*b*). The total real cost of *K* iterations is 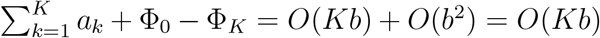, which means amortized cost of one iteration to be *O*(*b*).

Taken together the algorithm allows to find all cumulative enrichment scores *s_r_*(*π*[1..*k*]) in 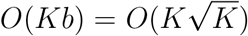 time. The straightforward implementation of calculating cumulative values from scratch would take *O*(*K*^2^ log *K*) time. Thus, we have improved the performance 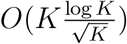 times.

#### 2.3.3 Implementation details

We also implemented an optimization so that the algorithm does not build convex hull from scratch for a changed block *c*, but only updates the changed points. This does not influence the asymptotic performance, but decreases the constant factor.

First, we start updating the convex hull from position *r_k_* and not from the start. To be able to do this, we have an array `prev` that for each gene *g* ∈ *π* stores the previous point on the convex hull if *g* were the last gene in the block. This actually is the same as the top of the stack in Graham algorithm and represent the algorithms state for any given point. As all points *h* to the left of *g* are not changed `prev`_*h*_ also remains unchanged and need not to be recalculated.

Second, we stop updating the hull, when we reach the point on the previous iteration convex hull. We can do this because every point to the left of *g* is rotated counterclockwise of any point to the right of *g*, which means that the first point on the convex hull right of *g* on (*k* – 1)-th iteration remains being a convex hull point at *k*-th iteration.

### 2.4 An algorithm for exact calculation of GSEA P-values for integer gene-level statistics

In this section we describe a polynomial algorithm to calculate GSEA P-value exactly, but only for the case when gene-level statistics are integer numbers: 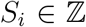. For simplicity we will consider a problem of calculating the following probability:

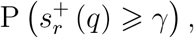

where *q* is a random gene set of size *k*. We also assume *γ* > 0.

Let denote the sum of *k* largest absolute values of gene ranks by *T*. The algorithm will be polynomial in terms of *N, k* and *T*.

#### 2.4.1 The basic algorithm

Let us consider a gene set *q* = {*q*_1_, *q*_2_,…, *q_k_*}. Recall the formula for *s*^+^(*q*):

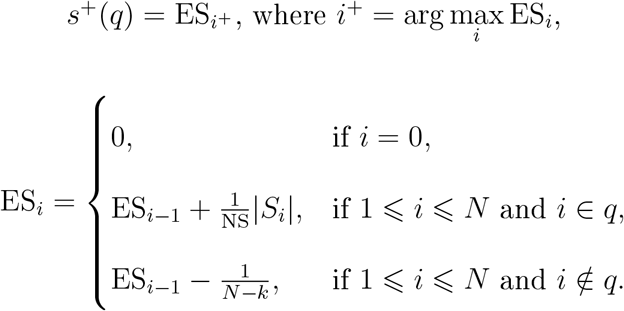

First, let rewrite the formula for ES_*i*_ in an equivalent fashion, grouping positive and negative summands:

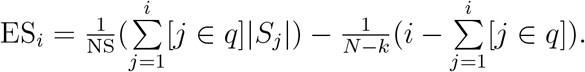

Then for calculating ES_*i*_ the following values are sufficient:

- *i*: the index of the current gene;
- 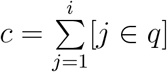: the number of genes included into the set *q* among genes 1..*i*;
- 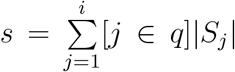: the sum of the absolute values of gene-level statistics for genes included in the set among genes 1..*i*;
- 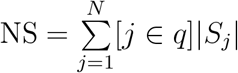: the sum of the absolute values of gene-level statistics for *all* genes in the set.

Knowing the values above, ES_*i*_ can be calculated as 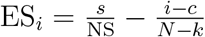.

Notice that NS can take only integer values from 0 to *T* (for a set of genes with the largest absolute values of gene-level statistics). Let us split the desired probability to a sum of independent probabilities based on the value of NS:

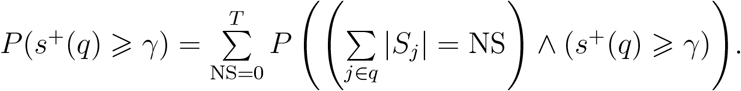

Our algorithm will be based on *dynamic programming*. For each possible value of NS we will process the genes one by one in increasing order of index and calculate an array *f*_NS_(*i, c, s*). The value *f*_NS_(*i, c, s*) will contain the probability for a uniformly random gene set *q*′ of *c* genes selected from genes 1..*i* to simultaneously have the following two properties:

1. the sum of the absolute values of gene-level statistics of genes from *q*′ is equal to *s*;
2. ES_*j*_ < *γ* holds for all *j* ⩽ *i*, where the values of ES are calculated for the gene set *q*′ but using the selected values of NS and *k*, not the ones calculated for the set *q*′. Suppose that we have calculated all values of *f*_NS_(*i, c, s*), then

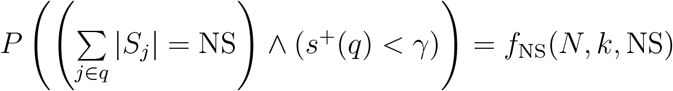

and

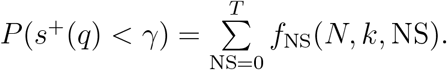 Finally, the sought probability is equal to:

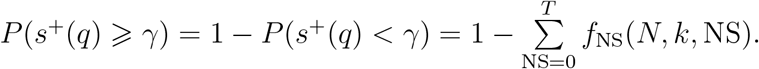 Let us find a formula for *f*_NS_(*i, c, s*). The base case of dynamic programming is *i* = 0 for all NS:

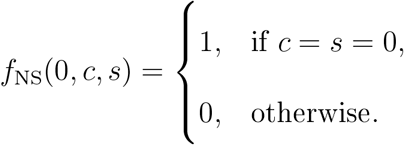 Suppose we want to calculate *f*_NS_(*i, c, s*) for some *i* > 0. First, calculate

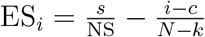

and compare it to *γ*. If ES_*i*_ ⩾ *γ*, then *f*_NS_(*i, c, s*) = 0 by definition.

Otherwise, condition “ES_*j*_ < *γ* holds for all *j* ⩽ *i*” can be simplified to “ES_*j*_ < *γ* holds for all *j* ⩽ *i* – 1”. This observation allows us to use values of *f* that have already been calculated. Consider two cases:

1. Gene *i* does not belong to the set *q*′. As *q*′ is a set of *c* genes chosen uniformly at random from *i* genes, this case happens with the probability 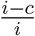. The conditional probability that such set satisfies the two necessary properties is *f*_NS_(*i* – 1, *c, s*). Indeed, any set of size *c* with the sum of absolute values of gene-level statistics values equal to *s*, chosen among genes 1..*i* – 1 and satisfying the conditions on ES, is a valid set chosen among genes 1..*i*. Similarly, if a set does not satisfy the condition on ES_*j*_ for some *j* ⩽ *i* – 1, this set should not be counted towards *f*_NS_(*i, c, s*) since obviously *j* ⩽ *i*.
2. Gene *i* belongs to the set. This case happens with the probability 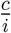. The probability that this set satisfies the necessary conditions is *f*_NS_(*i* − 1, *c* − 1, *s* − *S_i_*). Indeed, any set of size *c* − 1 with the sum of absolute values of gene-level statistics equal to *s* − *S_i_*, chosen among genes 1..*i* − 1 and satisfying the conditions on ES, can be extended with gene *i*, thus forming a set of size *c* satisfying both necessary properties. Similarly, if a set does not satisfy the condition on ES_*j*_ for some *j* ⩽ *i* − 1, adding gene *i* will not fix the situation. Then we can calculate *f*_NS_(*i, c, s*) using the law of total probability:

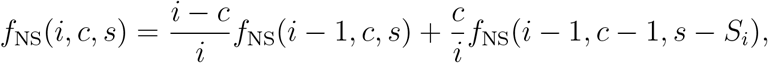

in the case when *i* > 0 and ES_*i*_ < *γ*. Putting all the cases together, we arrive to the final formula for *f*_NS_(*i, c, s*):

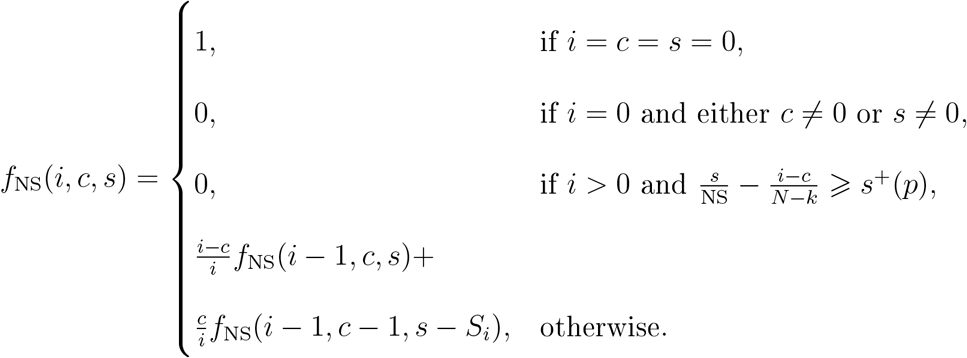 The overall complexity of the algorithm is *O*(*NkT*^2^). The values of *f* can be evaluated sequentially in increasing order of *i*. It is enough to evaluate *f*_NS_(*i, c, s*) for 0 ⩽ *i* ⩽ *N*, 0 ⩽ *c* ⩽ *k*, and 0 ⩽ *s* ⩽ NS ⩽ *T*. Each value of *f* can be evaluated in constant time.

#### 2.4.2 Optimizations and implementation details

While the algorithm described above is polynomial, a number of further optimizations are required to make execution on real size inputs feasible.

First, let note that the following property holds: *f*_NS_2__(*i, c, s*) ⩾ *f*_NS_1__(*i, c, s*) as long as NS_2_ ⩾ NS_1_. Indeed, ES values calculated using different values of NS are decreasing when NS is increased. That means all gene sets counted towards *f*_NS_1__(*i, c, s*) should also be counted towards *f*_NS_2__(*i, c, s*) if NS_2_ ⩾ NS_1_.

Following the observation above, instead of calculating values of *f*_NS_(*i, c, s*) we will consider the values *g*(*i, c, s, b*) = *f*_*b*+1_(*i, c, s*) − *f_b_*(*i, c, s*). These values will contain the probability of a random gene set *q* of size *k* selected uniformly from genes 1..*N* to satisfy simultaneously the following three properties:

1. set *q* contains exactly *c* genes from the genes 1..*i*.
2. the sum of the absolute values of gene-level statistics of the first *c* genes from *q* is equal to *s*;
3. ES_*j*_ < *γ* holds for all *j* ⩽ *i*, where the values of ES are calculated for the gene set *q* using NS = *b* +1 (and for all higher values of NS);
4. ES_*j*_ ⩾ *γ* holds for at least one *j* ⩽ *i*, where the values of ES are calculated for the gene set *q* using NS = *b* (and for all lower values of NS).

The sought probability can be calculated from values of *g* as follows:

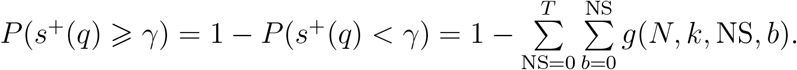

To calculate the values of *g* we will use the forward dynamic programming algorithm. In this algorithm we expand a tree of reachable dynamic programming states, starting from *g*(0, 0, 0, 0) which is equal to 1.

The states will be considered by “levels” in an increasing order of *i*. The values *g*(*i* + 1, *c, s, b*) from (*i* + 1)-th level are calculated based on level *i*. Note, that the sum of values on *i*-th level is always equal to 1.

To calculate all values from the (*i* + 1)-th level all non-zero values from the *i*-th level are considered sequentially. Let consider state (*i, c, s, b*) and let define *p* = (*k* – *c*)/(*N* – *i*) – the probability that gene *i* + 1 will be added to the set. The corresponding set *G*(*i, c, s, b*) can be divided into two groups.

1. The gene sets from *G*(*i, c, s, b*) that do not include gene *i* + 1. These gene sets are included into gene sets *G*(*i* + 1, *c, s, b*) on the level *i* + 1. Thus the corresponding probability *g*(*i, c, s, b*) · (1 – *p*) is added to the value of *g*(*i* + 1, *c, s, b*).
2. The gene sets from *G*(*i, c, s, b*) that do include gene *i* + 1. These gene sets are included into *G*(*i* + 1, *c* + 1, *s*′ = *s* + |*S*_*i*+1_|, *b*′) where *b*′ is an updated bound. To calculate *b*′ let note th at ES_*j*_ will be greater or equal to *γ* iff 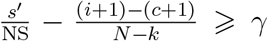 which is equivalent to 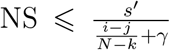. Thus 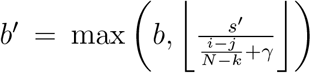 The probability that is added to *g*(*i* + 1, *c* + 1, *s*′, *b*′) is equal to *g*(*i, c, s, b*) · *p*.

While the asymptotic number of states remains to be *O*(*NkT*^2^) the forward dynamic programming allows to consider only “reachable” gene stats with *g*(*i, c, s, b*) > 0. In practice the number of reachable stats can be several orders of magnitude smaller then the total states.

Furthermore, for the algorithm we can consider only states with *g*(*i, c, s, b*) > *ε* to be reachable for some small value of *ε*. If we do not consider the unreachable states we would not be able to calculated the desired probability exactly. However, if we calculate the value of *δ* as a sum of all the skipped states values, the desired probability will be calculated with the absolute error no more than *δ*.

The algorithm implementation with few other optimizations is available at: https://github.com/ctlab/fgsea/blob/master/inst/exact/exact.cpp.

### 2.5 FGSEA-multilevel: an algorithm for calculation of arbitrarily low P-values using adaptive multilevel split Monte Carlo scheme

In this section we describe FGSEA-multilevel algorithm that can accurately estimate GSEA P-value for a pathway *p* of size *k* even when the true P-value is very small.

Let *γ* = *s_r_*(*p*) > 0 be the enrichment score of the query pathway *p* for which we want to calculate the following value:

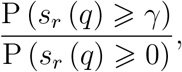

where *q* is a random gene set of size *k*. This probability can be rewritten as follows:

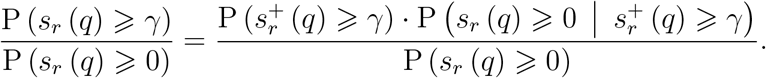

First, we focus on determining the probability 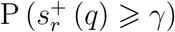. This probability can be extremely small, so using a naive sampling gives a bad estimation. We use the adaptive multilevel split Monte Carlo method [6] to solve this problem.

To estimate the probability 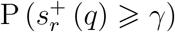 we split the enrichment scores into levels 0 = *l*_0_ < *l*_1_ < … < *l_t_* = *γ*. Then we can define the following probabilities:

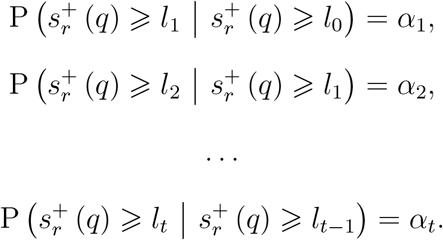

Now the probability 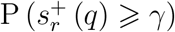 can be rewritten as 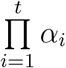.

To estimate *α_i_* we can draw a sample 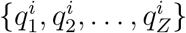 of size *Z* from a conditional distribution 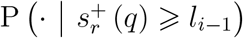. Then

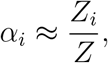

where *Z_i_* is the number of elements in the set 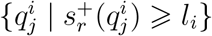.

Below we show how levels *l_i_* can be chosen and how to sample from the corresponding conditional distributions.

#### 2.5.1 Choosing the enrichment score levels

We propose to chose value for a level *l_i_* as a median of the enrichment scores for the 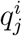 sample. For simplicity *Z* is required to be an odd number.

Then the procedure for estimating probability 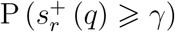 consists of repetition of the following steps:

1. On iteration *i* ⩾ 1 sample *Z* gene sets 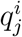 of size *k* from the distribution 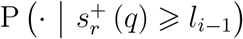.
2. Set the level 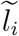 to be equal to the median of values 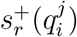.
3. If 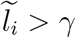 then stop the iterations and set *l_i_* = *γ* mid *t* = *i*, otherwise set 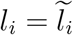.

As a result, by construction, *α_i_* ≈ 1/2 for 1 ⩽ *i* ⩽ *t* − 1. The value of can be approximated as *Z_t_*/*Z* (which is always ⩾ 1/2). Together we get the following expression for estimating the desired probability:

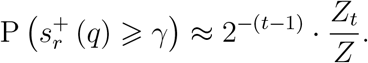

#### 2.5.2 The conditional sampling implementation

To generate a uniform sample 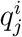 from the conditional distribution 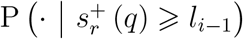 we use the Metropolis algorithm.

First, we generate a sample 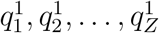 of size *Z* from the distribution 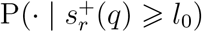. Since *l*_0_ = 0 and values of 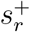 are always non-negative it can be done by generating a uniformly random subset of size *k* from the genes {1, 2,…, *N*}.

Now let consider a sample 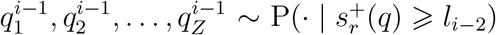 at a step *i* > 1. The sample can be sorted in an increasing order of enrichment score values: 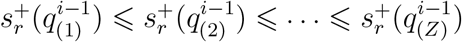. Let *d* = ⌈*Z*/2⌉. The level *l*_*i*−1_ is the median of the values 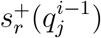 and, thus, is equal to 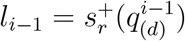.

Let first populate 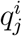 in the following way:

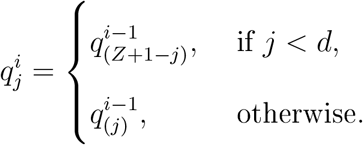

This gives us a sample from the conditional distribution 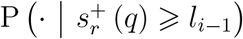, however it is not uniform.

To make the sample uniform we apply a number of the Metropolis algorithm iterations. On each iteration for each gene set 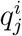 we apply the following steps:

1. Choose a random gene 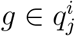.
2. Choose a random gene 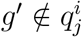.
3. Consider 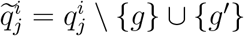. If 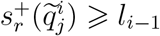 then we replace 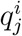 with 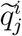.

The iterations are repeated until the total number of successful replacements becomes greater or equal to *k* · *Z*. In practice, this number of steps is enough to get a sufficiently uniform sample to obtain a good estimation of probability, without a significant increase in the running time of the algorithm.

#### 2.5.3 Estimating the P-value

In order to estimate the desired P-value we also need to calculate the probabilities P (*s_r_*(*q*) ⩾ 0) and 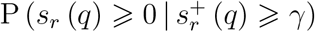.

To calculate the probability P (*s_r_*(*q*) ⩾ 0) we generate gene sets *q*_1_, *q*_2_,…, *q_Z_*′, where each sample *q_i_* is selected uniformly at random from all the subsets of size *k* from the set {1, 2,…, *N*}. The samples are generated until the number of samples *q_i_* with *s_r_*(*q_i_*) ⩾ 0 becomes equal to *Z*. Then the probability P (*s_r_*(*q*) ⩾ 0) is estimated as follows

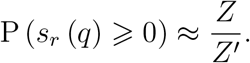

To determine the remaining probability 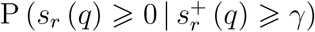 we calculate the number of gene sets in 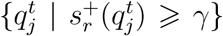 with value of the enrichment score function *s_r_* is greater than zero. After that, the probability can be estimated as follows:

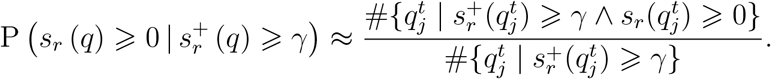

#### 2.5.4 Estimating log-probability

To properly estimate a logarithm of the desired probability let note that the *j*-th order statistic of a standard uniform sample of size *Z* is a random variable from the beta distribution Beta (*j, Z* + 1 − *j*). Therefore, we can use the properties of the beta distribution and make correct transition to the logarithm of probability. So for the median value of sample of odd size *Z* we have:

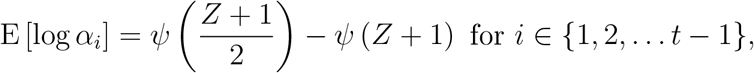

where *ψ* is digamma function. In the same way, we can calculate the expectation of the logarithm *α_t_*:

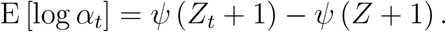

Then the logarithm of probability 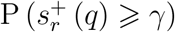 is estimated as

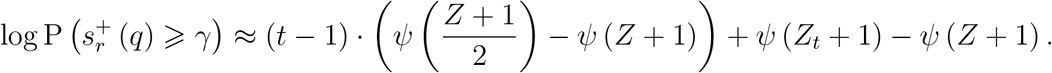

Similarly, we can estimate the variance of the estimates 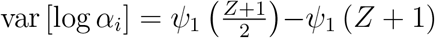, where *ψ*_1_ is trigamma function. From this we can approximate a standard error of our estimator as:

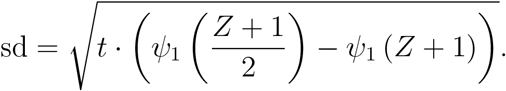

The same approach with digamma functions is used to calculate the logarithm of the probabilities 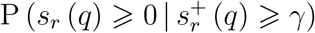 and P (*s_r_* (*q*) ⩾ 0).

#### 2.5.5 Optimizations and implementation details

The most demanding step of FGSEA-multilevel algorithm is to check whether the newly obtained gene set has the enrichments score value of at least *l_i_*. Importantly, this does not require full computation of enrichment score value. It is enough to show that there is at least one gene that has the running enrichment score value of at least *l_i_* or, in other words, that at least one point on the enrichment score graph is farther away from the diagonal (Fig. S1) than some value calculated from *l_i_*.

Similarly to FGSEA-simple algorithm, a square root decomposition is used to split all the genes into approximately 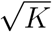 blocks. The block boundaries are determined automatically on each level based on the existing sample in such a way, that each block contains approximately 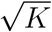 genes. In particular this decomposition enables adding and removing genes in 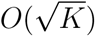 time while keeping the gene set sorted.

Unlike FGSEA-simple algorithm we will not maintain the convex hull but will apply a few heuristics to do the required check faster.

First we check a “candidate” point which has a high chance to be at the required distance from the diagonal. If it is so, we do not have to continue the check. The “candidate” gene is carried our from the previous iterations, as the point where the successful check has been interrupted.

Next we go block by block. At the beginning we construct a “rectangle” upper bound on the enrichment score value at the block, which can be obtained by moving all the genes of the block to its start. If this upper bound does not satisfy our criterion we can skip the block. Otherwise, we go gene by gene and calculate the enrichment score values until it reaches the required value or the end of the block is reached. In the former case the check is interrupted with a successful result.

#### 2.5.6 Comparison with the exact method

To compare FGSEA-multilevel and the exact method on the same dataset we used rounded values of the gene-level statistics from the example data (section 2.2) as input data for both algorithms. Both algorithms calculated the probability 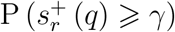.

The results of the algorithms for the pathways from the example data are shown on Fig 3c. The exact algorithm was run with *ε* = 10^−40^, all the probabilities were obtained with accuracy of at least six significant digits. For FGSEA-multilevel *Z* = 101 was used.

We also calculated empirical estimation errors and compared it to the theoretical ones (Fig 3d). For this we generated 100 independent estimates for a range of ES values (corresponding to P-values of 10^−4^ to 10^−100^, gene set sizes (from 15 to 250) and sample size (from 101 to 1001). The raw values are available in the Supplementary Table.

### 2.6 Filtering redundant pathways

In this section we describe an algorithm to filter redundant pathways from the results of FGSEA.

Let consider two pathways *p*_1_ and *p*_2_ that both have a significant GSEA P-value. There are two situations in which we will consider *p*_2_ to be non-redundant given *p*_1_:

1. If pathway *p*_2_ is enriched even if we do not consider the genes from *p*_1_ at all. Formally, we calculate GSEA P-value for gene set *p*_2_ \*p*_1_ and gene-level statistics vector *S*[*U* \*p*_1_] for all the genes except *p*_1_. If the P-value is less than a pre-defined threshold, then pathway *p*_2_ is considered as non-redundant given *p*_1_.
2. If pathway *p*_2_ is enriched even if we consider only genes from *p*_1_. Formally, we calculate GSEA P-value for gene set *p*_2_ ∩ *p*_1_ and gene-level statistics vector *S*[*p*_1_] for the genes from *p*_1_. Again, if the P-value is less than a pre-defined thresh old, then pathway *p*_2_ is considered as non-redundant given *p*_1_.

Otherwise pathway *p*_2_ is considered to be redundant.

The filtering procedure starts with a set of significantly enriched pathways *P_sig_* selected by the user: for example the pathways with GSEA P-values less than 0.01 after Benjamini-Hochberg correction, sorted by P-value. The output of the procedure is a list *P_main_* ⊂ *P_sig_* of pathways that are pairwise non-redundant. At the same time, all the other pathways *P_red_* = *P_sig_* \ *P_main_* are redundant given some pathway from *P_sig_*.

The procedure itself is similar to Sieve of Eratothenes algorithm. The pathways are considered one by one and some of them are marked as redundant. For a pathway *p* we first check if it is already marked as redundant, if yes, we go to the next pathway. Otherwise, we first run FGSEA-simple algorithm on a vector of statistics *S*[*U* \ *p*] and all the pathway currently not marked as redundant (including the ones that already have been considered, but excluding pathway *p*). Then, similarly, we run FGSEA-simple algorithm on a vector of statistics *S*[*p*]. Pathways that do not achieve non-redundant P-value threshold in both tests are marked as redundant.

---

**Figure S1:**
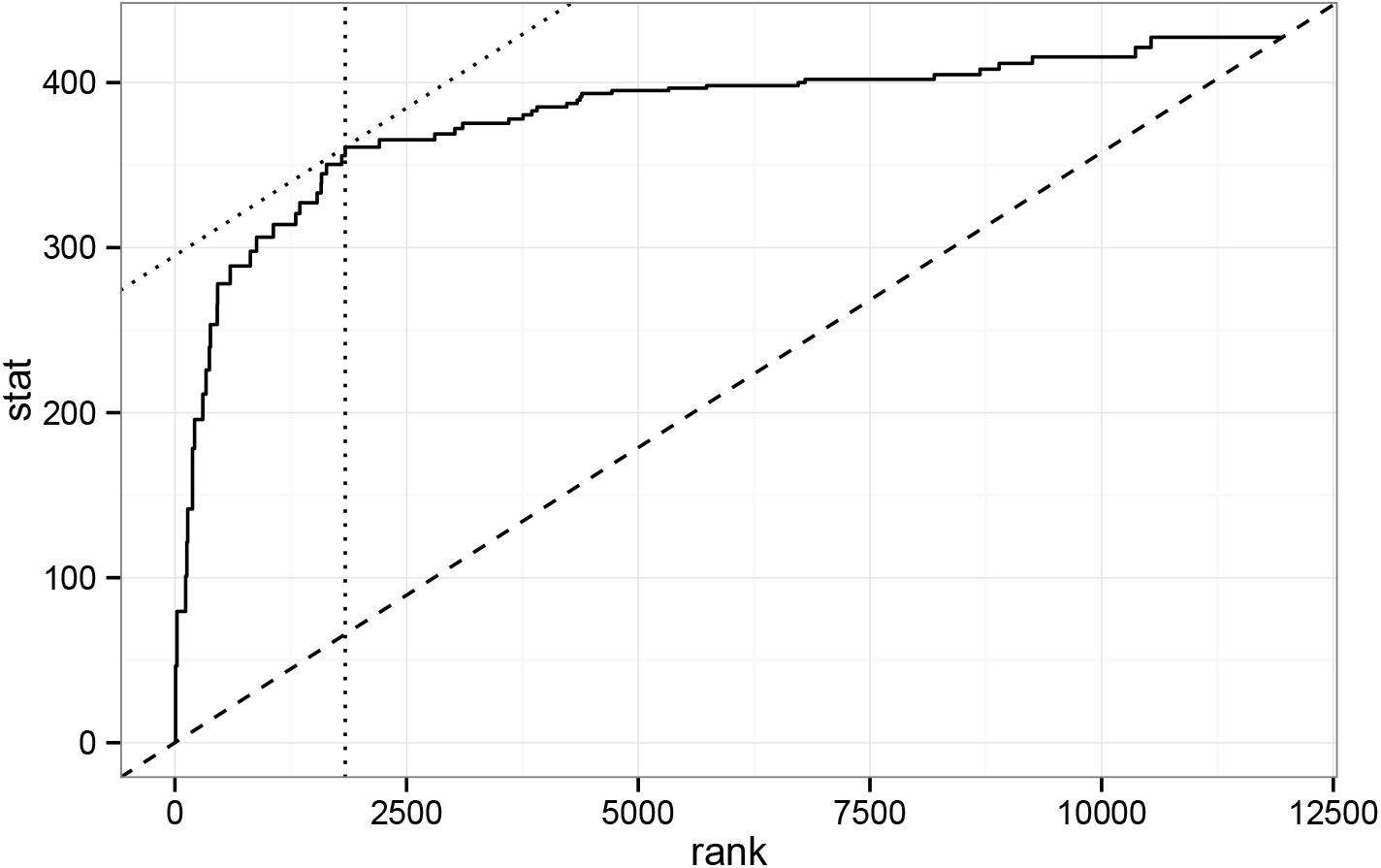
A graph that corresponds to a calculation of enrichment score. Each breakpoint on a graph corresponds to a gene present in the pathway. Dotted lines cross at a point which is the farthest up from a diagonal (dashed line). This point correspond to gene *i*^+^, where the maximal value of ES_*i*_ is reached.

**Figure S2:**
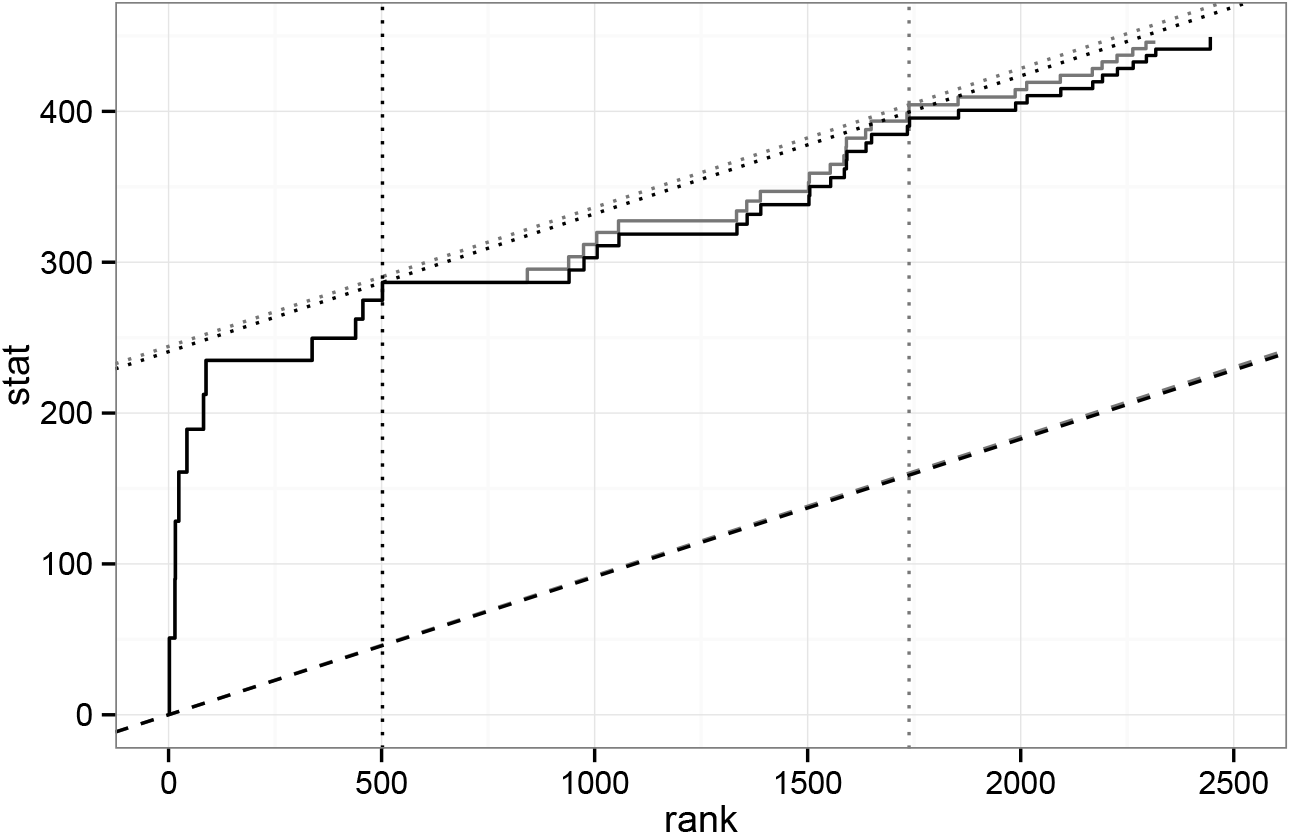
Update of an enrichment score graph when gene *π_k_* ≈ 800 is added. Only a fragment is shown. Black graph corresponds to a graph for gene set *π*[1..*k* − 1], gray graph corresponds to *π*[1..*k*], A part of the graph to the left of *x* = *x_r_k__* does not change and the other part is shifted to the top-left corner. The diagonal ((*x*_0_, *y*_0_), (*x_N_, y_N_*)) is rotated counterclockwise.

## Notes

https://github.com/ctlab/fgsea/

### Competing Interest Statement

The authors have declared no competing interest.

### Summary of Updates

A systematic analysis of GSEA performance on 605 datasets has been added.

